# Activation of 2,4-diaminoquinazoline in *Mycobacterium tuberculosis* by Rv3161c, a putative dioxygenase

**DOI:** 10.1101/375519

**Authors:** Eduard Melief, Shilah A. Bonnett, Edison S. Zuniga, Tanya Parish

**Affiliations:** TB Discovery Research, Infectious Disease Research Institute, Seattle, WA, 98102 USA

**Author notes:** Address correspondence to Tanya Parish.

**Keywords:** Diaminoquinazoline, *Mycobacterium tuberculosis*, anti-tubercular, prodrug, activation, monooxygenases, dioxygenases

## Abstract

The diaminoquinazoline series has good potency against *Mycobacterium tuberculosis*. Resistant isolates have mutations in Rv3161c, a putative dioxygenase. We carried out metabolite analysis on wild-type and an Rv3161c mutant strain after exposure to a diaminoquinazoline. The parental compound was found in intracellular extracts from the mutant, but not the wild-type. A metabolite consistent with a mono-hydroxylated form was identified in the wild-type. These data support the hypothesis that Rv3161c metabolizes diaminoquinazolines in *M. tuberculosis*.

---

The diaminoquinazoline scaffold has been utilized in the generation of various tool and lead like compounds for anticancer and antimalarial drug discovery programs (1, 2). A high throughput screening campaign led to the discovery of a series of diaminoquinazoline (DAQ) compounds that was active against *Mycobacterium tuberculosis* with minimum inhibitory concentrations (MIC) in the sub-micromolar range (3). We carried out a structure activity relationship analysis to evaluate the potential of the DAQ series and identified a number of analogs with improved anti-tubercular activity and good exposure in rat pharmacokinetic studies (4). DAQ compounds had bactericidal activity against replicating and non-replicating bacteria (4). The target for the DAQ series is not known, but DAQ-resistant isolates have mutations in Rv3161c, a putative dioxygenase (4). Since Rv3161c is not essential, and we predict that the mutation would lead to reduced or lower activity, it is unlikely that this is the intracellular target of the series.

Rv3161c is highly induced in *M. tuberculosis* treated with benzene-containing compounds such as thioridazine, SRI#967, SRI#9190 and triclosan (5–7). However, triclosan was equally active against wild-type and Rv3161c deletion strains of *M. tuberculosis*. In addition, strains which over-expressed Rv3161c did not demonstrate triclosan resistance, suggesting it is not involved in mediating resistance (5).

We hypothesize that DAQ compounds are prodrugs activated by Rv3161c. In order to assess this hypothesis we wanted to determine if DAQ molecules were transformed after uptake by *M. tuberculosis*. In our previous work, we had only determined the MIC on solid medium using the serial proportion method (8). For compound IDR-0010006, the MIC_99_ for wild-type was 6.25 μM, but for the resistant mutants was 25 μM (a 4-fold shift) (4). In order to run metabolite identification studies, we determined the MICs for the wild-type and resistant strains in Middlebrook 7H9 liquid medium supplemented with 10% v/v OADC (oleic acid, albumin, dextrose, catalase) and 0.05% w/v Tween 80. Compounds were tested as a 10-point, 2-fold serial dilution in 96-well plates as described (9)(8). Growth was measured after 5 days at 37°C and the % growth calculated with respect to controls. Curves were fitted using the Levenberg-Marquardt algorithm and IC_90_ calculated as the concentration of compound required to inhibit growth by 90%. We determined IC_90_s for IDR-0010006 and IDR-0258237 against wild-type and the mutants strain containing the all Rv3161c_C115W_ allele (Table 1). Both compounds were 2.5-fold less active against the mutant strain as compared to wild-type, which is comparable to the shift seen on solid medium.

**Table 1.**
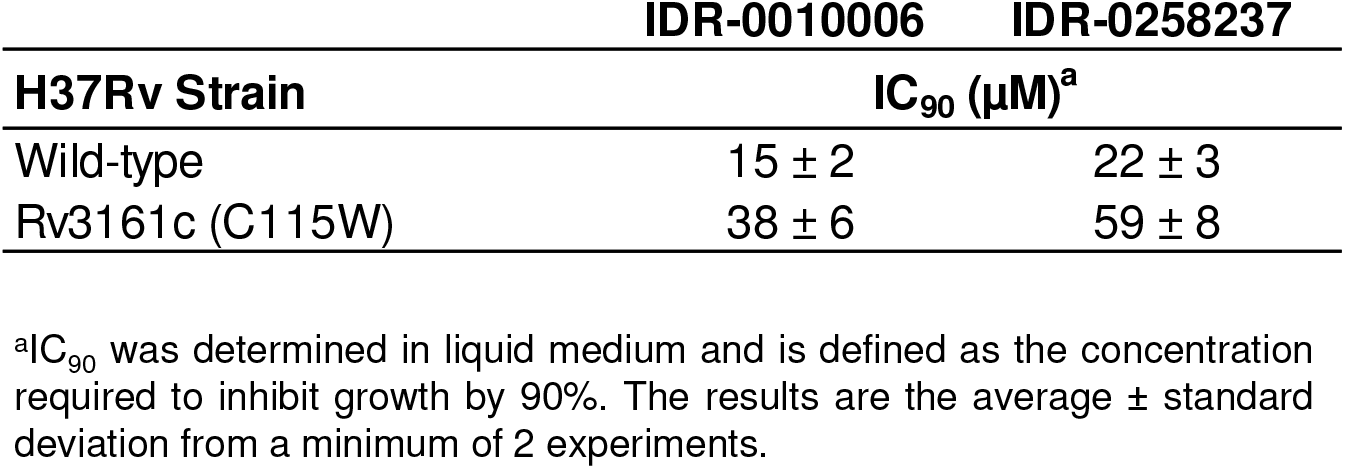
Activity of DAQ compounds

|  | IDR-0010006 | IDR-0258237 |
| --- | --- | --- |
| H37Rv Strain | IC<sub>90</sub> (μM)<sup>a</sup> |  |
| Wild-type | 15 ± 2 | 22 ± 3 |
| Rv3161c (C115W) | 38 ± 6 | 59 ± 8 |

<sup>a</sup>IC<sub>90</sub> was determined in liquid medium and is defined as the concentration required to inhibit growth by 90%. The results are the average ± standard deviation from a minimum of 2 experiments.

Once we had established liquid IC_90_, we carried out metabolite analysis using compound IDR-0258237. We first established the settings required to detect the parent compound. LC-MS was carried out on using an Agilent 1100 HPLC and G1956B LC/MSD SL mass spectrometer set in positive mode with simultaneous UV-vis detection at 214 and 254 nm. Buffer A was 0.05% formic acid in water and Buffer B was 0.05% formic acid in acetonitrile. The following solvent gradient was used (time, % Buffer B): 0 min, 10%; 30 min, 6.3 min, 95%; 8.1 min, 95%; 8.4 min, 10%; and 10.5 min, 10%. Under these conditions, compound IDR-0258237 eluted at ~4.5 min with a m/z value of 336, which is consistent with the calculated exact mass ([M+H]^+^) of the parent compound (Fig 1A).

**Fig. 1.**
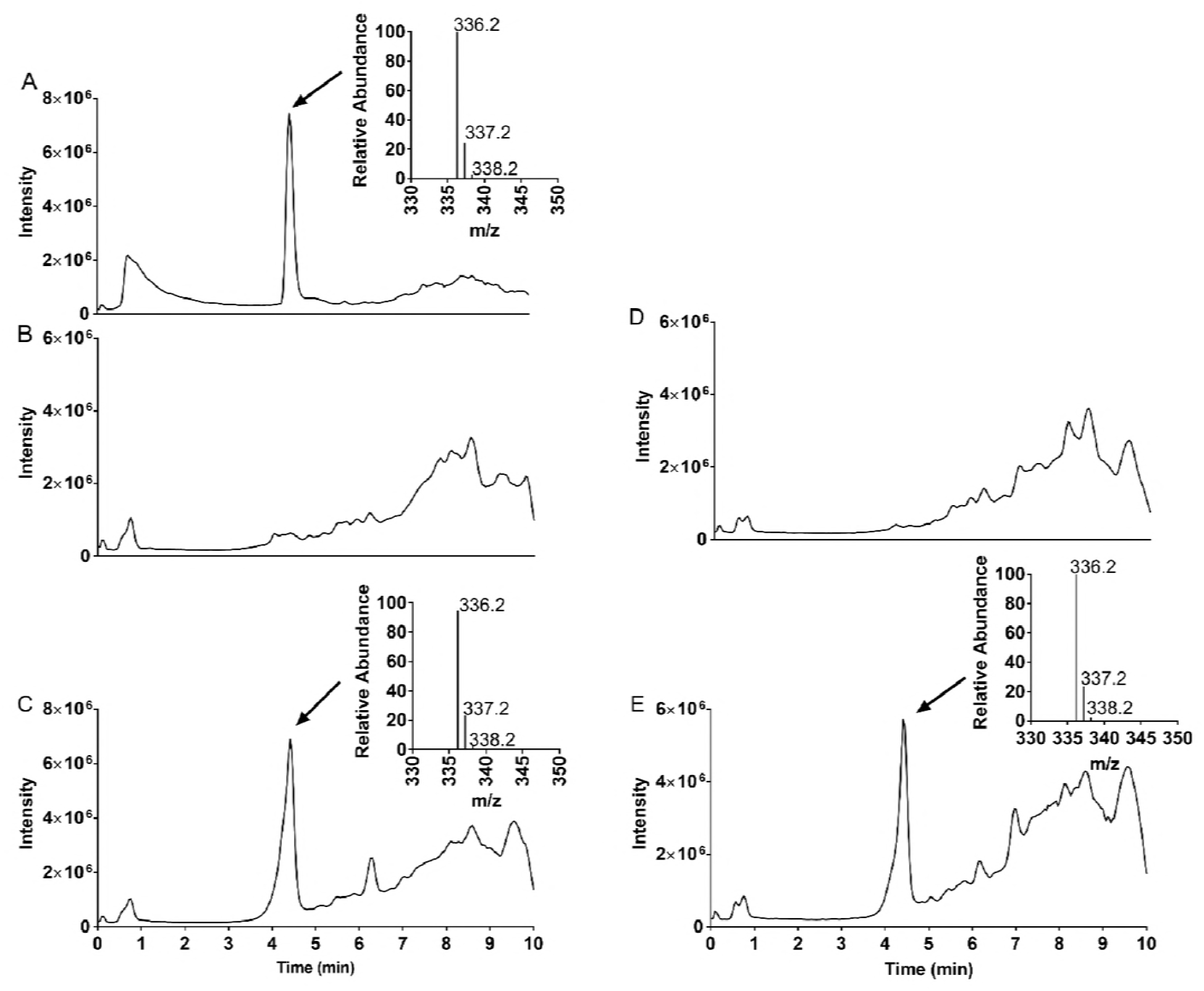
Metabolite analysis. (A) Total Ion Chromatogram for IDR-0258237. *M. tuberculosis* was treated with 20 μM of IDR-0258237 for 24 h (B) and (C) or for 48 h (D) and (E). Extracts were subjected to LC-MS. (B) and (D) wild-type strain. (C) and (E) Rv3161c mutant strain. The inset shows the m/z peaks associated with the parent ion.

Next, we treated wild-type and mutant strains with compound IDR-0258237. *M. tuberculosis* cultures were grown in 100 mL of 7H9-Tw-OADC in 450 cm^2^ roller bottles at 100 rpm for 5 days at 37°C.Compound IDR-0258237 was added at 20 μM for 24 h or 48h; DMSO was used as a negative control. Cells were harvested by centrifugation, extracted with an equal volume of 1:1 chloroform-methanol and incubated at 55°C for 30 min. Samples were re-extracted with 1:1 chloroform-methanol and the two extracts pooled. The volume was adjusted to 5 mL with 1:1 chloroform-methanol and refluxed for 16-24 hr. Pellets were extracted with chloroform, dried, resuspended in 1:1 acetonitrile:water and subjected to LC-MS analysis (Fig 1 and Fig 2).

**Fig. 2.**
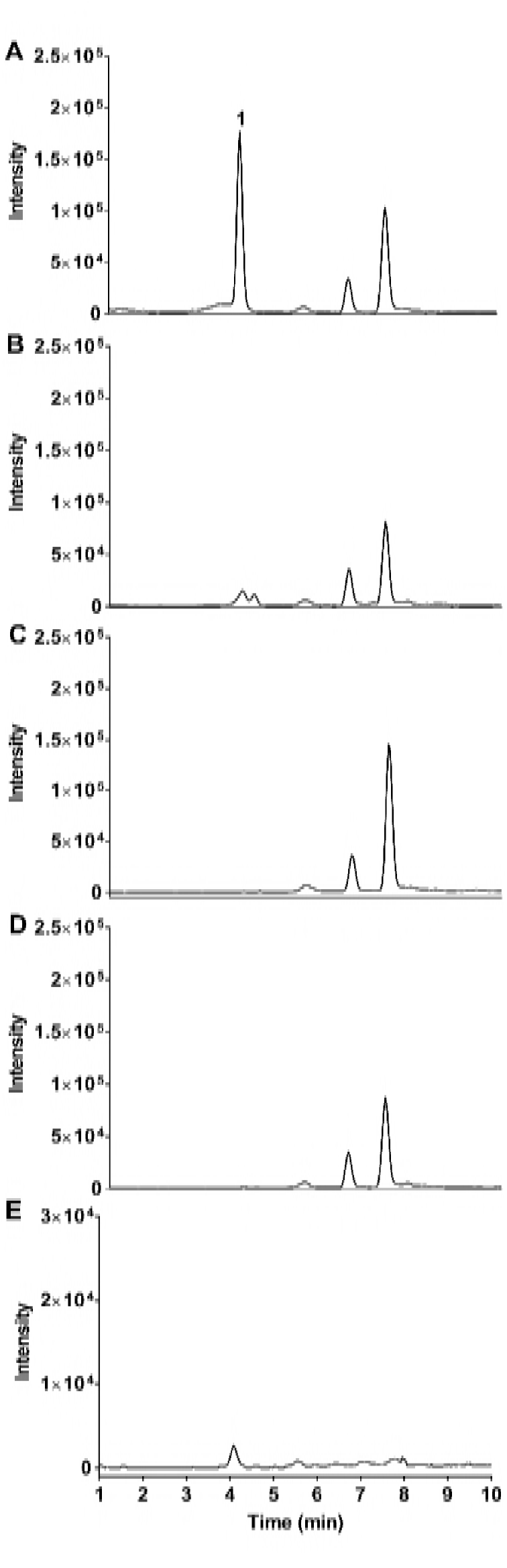
Extracted-Ion Chromatogram for m/z 354. (A) IDR-0258237-treated wild-type strain (B) IDR-0258237-treated Rv3161c mutant strain. (C) DMSO-treated wild-type strain. (E) DMSO-treated Rv3161c mutant strain. (E) Compound only.

In the wild-type strain, we did not see a peak corresponding to IDR-0258237 in the total ion chromatogram (TIC) (Fig 1B). In contrast, a peak consistent with IDR-0258237 was observed in extracts from the mutant strain (Fig 1B). Similar results were observed after 48 h of incubation (Fig 1D and 1E).

Rv3161c is a putative dioxygenase, which could catalyze the incorporation of two oxygen atoms into the substrate. Alternatively, it could function as a monooxygenase and catalyze the introduction of a single oxygen atom. We extracted the ion masses of all potential mono-hydroxylated and di-hydroxylated products from the TIC from all samples (Fig 2). An ion (**1**) with a retention time of ~4.1 min and m/z of 354, was observed in the wild-type extracts, but not in any of the controls or in the Rv3161c mutant extracts (Fig. 2). The MS properties of this ion are consistent with the addition of one water molecule across the heteroaromatic ring system resulting in the mono-hydroxylation of N1 (mono-hydroxylated DAQ) (Fig. 3). Alternatively, an epoxidation will also give the observed ion with an exact mass of 354. We did not detect any other mono-hydroxylated, di-hydroxylated or cleaved aromatic ring derivatives. LC/MS-MS experiments would be needed for metabolite ID and confirmation. These results are consistent with Rv3161c catalyzing the modification of DAQ compounds into a metabolite that is active against *M. tuberculosis*.

**Fig. 3.**
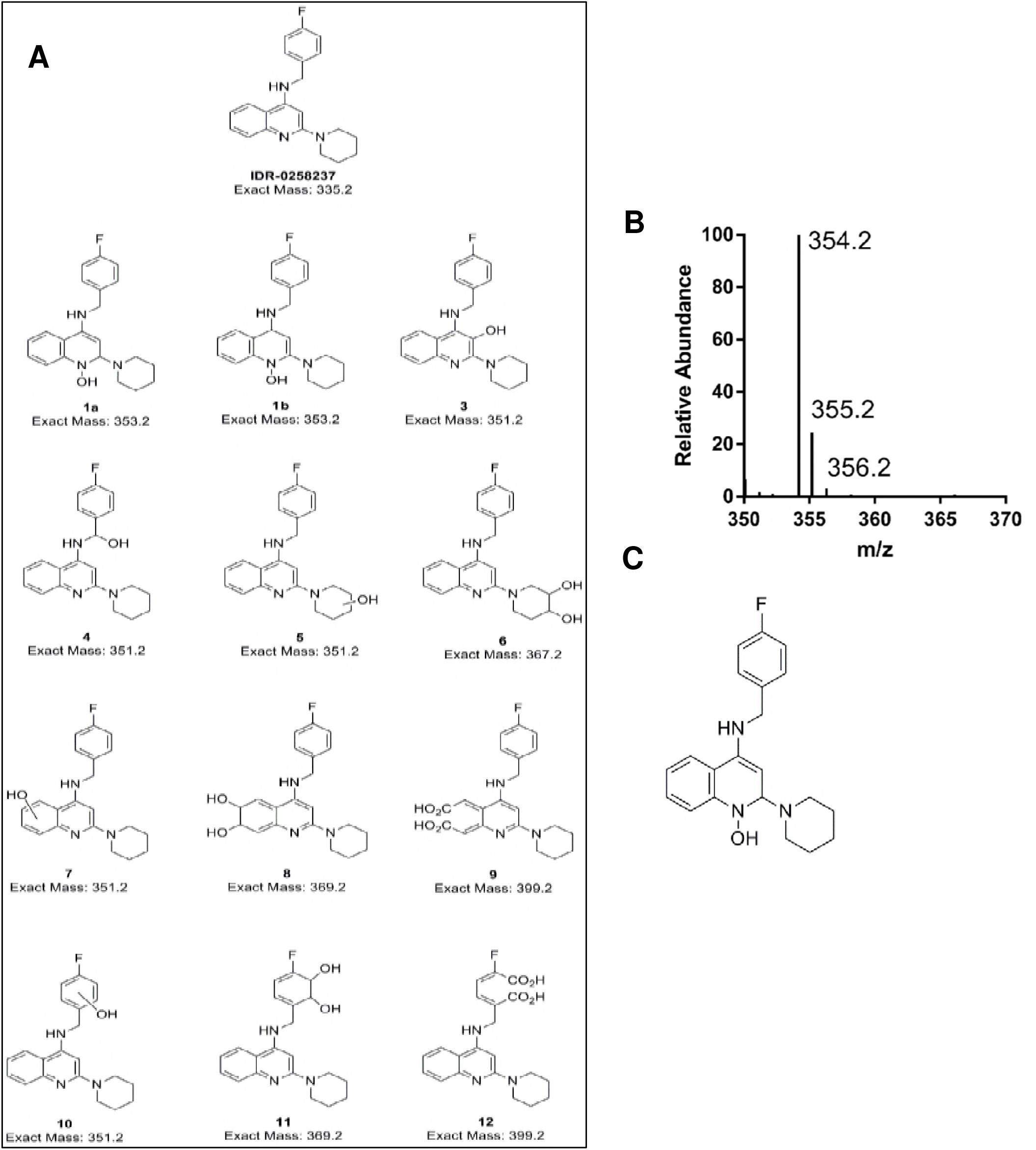
Identification of metabolites. (A) All possible metabolites. (B) MS spectra of the peak 1 detected in the extracted ion chromatogram. (C) Proposed structure of the active metabolite (m/z 354.2).

Mutation of Rv3161c only led to low level resistance, although the mutant strain did not appear to metabolize the DAQ compound to any detectable extent. This suggests that both the parent molecule and the metabolite are active, but that the metabolite has greater activity against the unknown target. We attempted to synthesize analogs incorporating the predicted hydroxylation, but we were not successful. Therefore, a full characterization of the metabolite(s) and its activity would require a large scale purification directly from *M. tuberculosis*.

In conclusion, we have determined that a member of the DAQ series is metabolized by wild-type *M. tuberculosis*, but not by a strain containing a mutation in Rv3161c. These data support our hypothesis that the DAQ compounds are biotransformed to more active compounds within the bacterial cell.

## Funding

This work was funded in part by Eli Lilly and Company in support of the mission of the Lilly TB Drug Discovery Initiative and by NIAID of the National Institutes of Health under award number R01AI099188.

